# Molecular bases for the constitutive photomorphogenic phenotypes in *Arabidopsis*

**DOI:** 10.1101/388157

**Authors:** Vinh Ngoc Pham, Xiaosa Xu, Enamul Huq

**Affiliations:** Department of Molecular Biosciences and The Institute for Cellular and Molecular Biology, The University of Texas at Austin, Austin, TX 78712

**Keywords:** E3 Ubiquitin ligase, Phytochrome Interacting Factors, photomorphogenesis, 26S Proteasome degradation, Skotomorphogenesis, Arabidopsis

## Abstract

Strikingly similar morphological and gene expression phenotypes among *cop1, spaQ* and *pifQ* mutants suggest that the *cop* phenotype of the *cop1* and *spaQ* mutants might be due in part to a reduced level of PIFs

**Abstract:** The transition from skotomorphogenesis to photomorphogenesis is regulated in part by COP1/SPA complex and PIFs in *Arabidopsis.* The constitutive photomorphogenic (cop) phenotypes of the *cop1* and *spaQ* mutants were shown to be due to a high abundance of the positively acting transcription factors. Here we show that the four major PIF proteins are unstable in *cop1* mutant, and an overexpression of *P1F1, P1F3, P1F4* and *P1F5* suppresses the *cop1* phenotypes in the dark. A comparison of the transcriptome data among *cop1, spaQ* and *pifQ* reveals remarkably overlapping gene expression profiles with a preferential regulation of the PIF direct target genes. Additionally, HFR1 strongly inhibits the *in vivo* binding and transcriptional activation activity of PIF1 in the dark. Taken together, these data suggest that the cop phenotypes of the *cop1* and *spaQ* mutants might be due to a combination of the reduced level of PIFs, increased level of the positive factors (e.g., HY5/HFR1 and others), and the HFR1-mediated inhibition of PIF targeted gene expression in the dark.

## Introduction

Plants are evolved with contrasting developmental programs for successful establishment of young postgerminating seedlings early in their life cycle. In darkness, plants undergo skotomorphogenesis defined by elongated hypocotyls, an apical hook and closed cotyledons. This developmental program is suited for protection of the apical region during rapid emergence of the seedlings through the soil surface. Once the seedlings are exposed to ambient light, they undergo photomorphogenesis defined by short hypocotyls, an absence of apical hook, open, expanded and green cotyledons. This growth pattern allows seedling body plan formation for maximal light capture and autotrophic growth (Gommers and Monte, 2018). Photomorphogenesis has been proposed to be the default pathway for plant development as a series of mutants displaying constitutive photomorphogenic (cop) phenotypes in the dark has been described (Xu, *et al,* 2015). These include 11 loci encoding the *CONSTITUTIVE PHOTOMORPHOGENIC1/DE-ETIOLATED1/FUSCA (COP/DET/FUS)* genes (Lau and Deng, 2012), four loci encoding *SUPPRESSOR OF PHYA-105 (SPA1-SPA4)* (Laubinger, *et al*, 2004) and a small family of basic helix loop helix (bHLH) transcription factor genes called the *PHYTOCHROME INTERACTING FACTORs (PIF1-PIF8)* (Leivar and Quail, 2011; Pham, *et al,* 2018b). These genes encode repressor proteins that act additively and/or synergistically to prevent precocious germination and seedling establishment in the dark.

Among these repressors, COP1 functions as an E3 ubiquitin ligase in association with SPA1-SPA4 targeting a variety of substrates including the positively acting transcription factors (e.g., HY5/HFR1/LAF1 and others) in light signaling pathways for Ubiquitin/26S proteasome-mediated degradation (Hardtke, *et al,* 2000; Hoecker, 2017; Jang, *et al,* 2005; Osterlund, *et al,* 2000; Seo, *et al,* 2003; Yang, *et al,* 2005b). COP1, SPAs and CUL4 form CUL4^COP1-SPA^ E3 ubiquitin ligase complexes that target positively acting factors in the dark (Zhu, *et al*, 2008). Consistently, *cop1, spaQ* and *cul4cs* (co-suppressed) lines display constitutive photomorphogenic (cop) phenotypes in the dark. In addition, another complex called the COP9 signalosome (CSN) consists of eight distinct subunits (CSN1-CSN8), which is involved in deconjugation of the NEDD8/RUB1 to the CULLIN RING ligases (CRLs) (Lau and Deng, 2012; Serino and Deng, 2003). Mutation in any of these subunits also display cop phenotypes in the dark.

DET1 is a nuclear protein, which binds to the N-terminal tail of histone H2B and regulates cell-type-specific expression of light-regulated genes (Benvenuto, *et al,* 2002; Pepper, *et al,* 1994). DET1 also promotes skotomorphogenesis in part by stabilizing PIFs in the dark (Dong, *et al,* 2014). In addition, DET1 suppresses seed germination by destabilizing HFR1 and stabilizing PIF1 (Shi, *et al,* 2015). DET1 also interacts with COP10 and DAMAGED DNA BINDING PROTEIN 1 (DDB1) to form CUL4^CDD^ complex that represses photomorphogenesis in the dark in part by degrading positively acting transcription factors (Chen, *et al,* 2006; Schroeder, *et al,* 2002).

PIFs belong to the basic helix-loop-helix (bHLH) family of transcription factors that repress photomorphogenesis in the dark by promoting skotomorphogenic development. There are eight PIFs (PIF1-PIF8) in Arabidopsis with a high degree of sequence similarity (Pham, Kathare and Huq, 2018b). However, individual *pif* mutants display distinct phenotypes that are especially pronounced in the four major *pif* mutants *(pif1, pif3, pif4* and *pif5* collectively called PIF quartet). For example, *pif1* seeds germinate under red and far-red light as well as in darkness (Oh, *et al,* 2004; Shen, *et al,* 2005), suggesting PIF1 is a repressor of light-induced seed germination. Both *pif1* and *pif3* mutants have more chlorophyll and carotenoids compared to wild type during the transition from dark to light (Huq, *et al,* 2004; Moon, *et al,* 2008; Stephenson, *et al,* 2009; Toledo-Ortíz, *et al,* 2010), suggesting that PIF1 and PIF3 suppress the biosynthesis of these pigments. *pif3, pif4* and *pif5* mutants display hypersensitive phenotypes in response to red light, in part by inducing co-degradation of these PIFs with phyB (Huq and Quail, 2002; Khanna, *et al,* 2007; Monte, *et al,* 2004; Zhu and Huq, 2014). In this process, multiple kinases (e.g., PPKs) and E3 Ubiquitin Ligases (e.g., CUL3^LRB^) participate in inducing the co-degradation of PIFs and phyB in response to light (Ni, *et al,* 2017; Ni, *et al,* 2014; Pham, Kathare and Huq, 2018b). Thus, phyB is more abundant in these mutants resulting in hypersensitive phenotypes under red light. In addition, other kinases (e.g., PPKs. BIN2 and CK2) and E3 Ubiquitin Ligases (e.g., CUL1^EBF1/2^, CUL3^BOP^ and CUL4^COP1-SPA^) induce the degradation of PIFs in response to light in a phytochrome-dependent manner to promote photomorphogenesis (Bernardo-García, *et al,* 2014; Bu, *et al,* 2011; Dong, *et al*, 2017; Ni, Xu, González-Grandío, *et al,* 2017; Pham, Kathare and Huq, 2018b; Zhang, *et al,* 2017). Strikingly, the quadruple mutant of the PIF quartet, *pifQ (pif1 pif3 pif4 pif5)* displays constitutive photomorphogenesis in the dark (Leivar, *et al,* 2008; Shin, *et al,* 2009), suggesting that these PIFs repress photomorphogenesis in the dark. They do so by regulating gene expression directly and indirectly in an individual to a shared manner (Pfeiffer, *et al,* 2014; Pham, Kathare and Huq, 2018b).

The cop phenotypes of the *cop1* and *spaQ* mutants were thought to be primarily due to a high abundance of the positively acting transcription factors (e.g., HY5/HFR1/LAF1 and others) in the dark (Hoecker, 2017). However, a few reports showed that PIFs are less abundant in *cop1* (Bauer, *et al,* 2004; Shen, *et al,* 2008; Xu, *et al,* 2017; Zhu, *et al,* 2015) and also to a lesser extent in the *spaQ* mutants (Leivar, Monte, Oka, *et al,* 2008; Ni, Xu, Tepperman, *et al,* 2014), suggesting that the instability of PIFs might contribute to the cop phenotypes of these mutants. Here we show that the gene expression signature of *cop1* and *spaQ* overlaps with *pifQ* in the dark with a preferential targeting of PIF direct target genes, suggesting that the cop phenotype of the *cop1* and *spaQ* is partly due to a reduced level of PIFs in these backgrounds. In addition, we also show that the positively acting transcription factor HFR1 strongly suppresses PIF1 function by sequestration; thereby, promoting the cop phenotypes of the *cop1* and *spaQ* in the dark.

## Results

### COP1 and SPA positively regulate PIF protein level in darkness

The constitutive photomorphogenic (cop) phenotypes of the *cop1-4, spaQ* and *pifQ* have been previously described (Figure 1A) (Deng, *et al,* 1992; Laubinger, Fittinghoff and Hoecker, 2004; Leivar, Monte, Oka, *et al,* 2008; Shin, Kim, Kang, *et al,* 2009). To examine if the cop phenotype of the *pifQ* is due to a reduction in the COP1 level, we performed immunoblot using an anti-COP1 antibody for four-day-old dark-grown seedlings of wild type, *cop1-4, spaQ* and *pifQ.* Results show that the COP1 level in *pifQ* and *spaQ* is similar to that in the wild type seedlings (Figure 1B), suggesting that the *pifQ* phenotype is not due to a reduction in the COP1 level.

**Figure 1:**
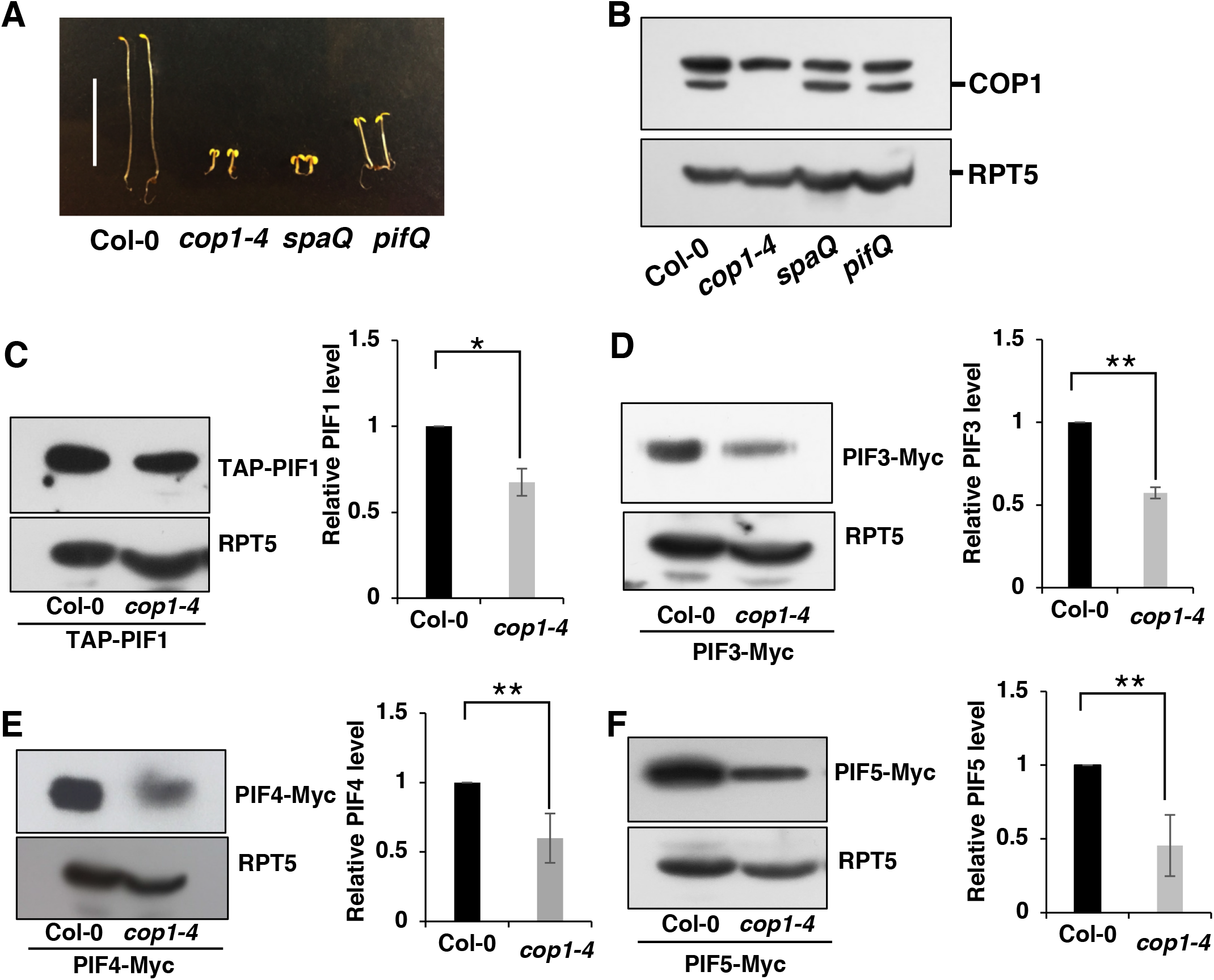
COP1 positively regulates PIF protein level in darkness. (A) Visible constitutive photomorphogenic phenotypes of 4-day-old dark grown seedlings. Bar = 10mm. (B) Immunoblot showing COP1 endogenous protein level in wild-type (Col-0), cop1-4, spaQ and pifQ dark-grown seedlings. Total protein was extracted from 4-day old dark-grown seedlings, separated on 8% SDS-PAGE gel, probed with anti-COP1 and anti-RPT5 antibody. (C-F) Immunoblots showing PIFs protein level and quantitative graphs. The overexpression plants TAP-PIF1, PIF3-Myc, PIF4-Myc and PIF5-Myc in Col-0 and cop1-4 backgrounds were grown under the condition described above. Total protein was separated in 6.5% SDS-PAGE gel and probed with anti-Myc and anti-RPT5 antibodies. PIFs protein level were quantified from three biological replicates (n=3) and normalized with RPT5 levels. The PIF protein level in Col-0 was set as 1. * indicates P < 0.05, ** indicates P < 0.01.

Previously, we and others have shown that the PIF levels in the *cop1* and *spaQ* mutants are reduced compared to wild type in the dark (Bauer, Viczian, Kircher, *et al,* 2004; Leivar, Monte, Oka, *et al,* 2008; Zhu, Bu, Xu, *et al,* 2015). To systematically analyze PIF levels without the transcriptional regulation in these mutants, we crossed the overexpression lines of tagged *PIF1, PIF3, PIF4* and *PIF5* using the constitutively active 35S promoter in the *cop1-4,* and TAP-PIF1 and PIF5-Myc in *spaQ* mutant backgrounds, and performed immunoblots for protein levels. Results show that all four PIF levels are reduced in the *cop1-4* as previously shown (Figure 1C-F). Both TAP-PIF1 and PIF5-Myc levels are reduced in the *spaQ* compared to wild type (Figure S1). Thus, the cop phenotype of the *cop1-4* and *spaQ* might be in part due to a reduction in the PIF levels in these backgrounds.

We also examined the transcript levels of the native *PIFs* in the *cop1-4* and *spaQ* mutants using quantitative RT-PCR assays (Figure S2). The transcript levels of *PIF1, PIF3* and *PIF4* are similar between Col-0 and *cop1-4,* except the transcript level of *PIF5* is strongly increased in *cop1-4.* In *spaQ* mutant, the transcript levels of *PIF1, PIF3* and *PIF4* were slightly lower, while *PIF5* transcript is slightly higher compared to wild type seedlings. These data illustrate that COP1 and SPA proteins positively regulate PIF protein levels in darkness possibly by destabilizing HFR1, as HFR1 has been shown to induce degradation of PIF1 by heterodimerization (Xu, Kathare, Pham, *et al,* 2017).

### PIFs are degraded in the *cop1-4* and *spaQ* mutants through the 26S proteasome

PIFs are known to be stable in the dark and undergo degradation in light through the 26S proteasome pathway (Pham, Kathare and Huq, 2018b). However, a recent study showed that PIF1 is also degraded in the dark by direct heterodimerization with HFR1 (Xu, Kathare, Pham, *et al,* 2017). To test whether the degradation of other PIFs in the *cop1-4* and *spaQ* backgrounds in the dark is also through the 26S proteasome pathway, we treated the dark-grown seedlings with the proteasome inhibitor (Bortezomib) for 4 hours and then extracted total protein for immunoblots. Results show that the proteasome inhibitor prevents the degradation of all four PIFs in the *cop1-4* background (Figure 2). Both TAP-PIF1 and PIF5-Myc are also stabilized in the *spaQ* background upon Bortezomib treatment (Figure S1). These data suggest that COP1/SPA complex stabilizes PIFs in the dark presumably by destabilizing HFR1.

**Figure 2:**
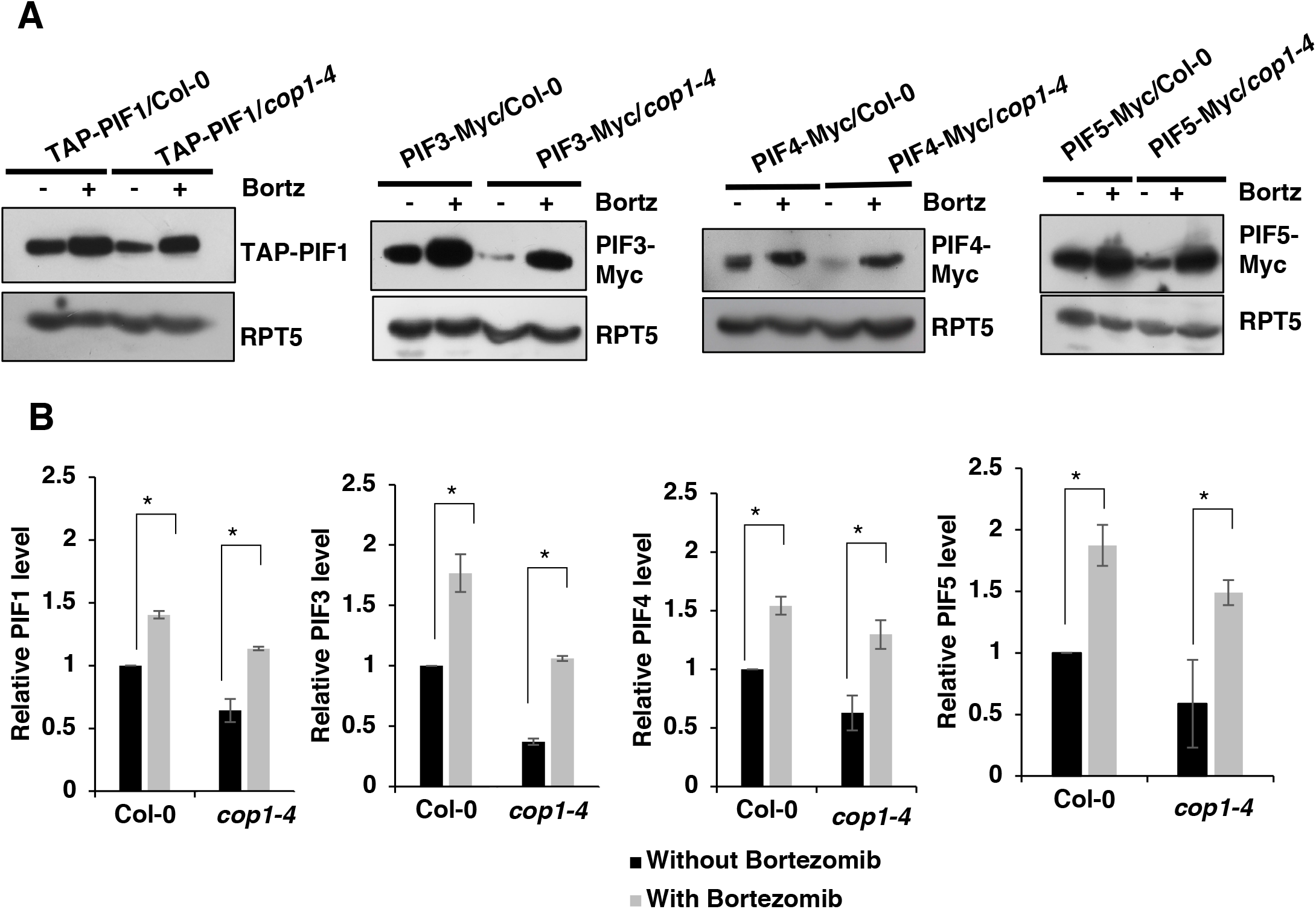
Instability of PIFs in the dark in *cop1* backgrounds is 26S-proteasome dependent. (A) PIF protein levels in 4-day-old dark-grown wild-type and cop1-4 seedlings with and without treatment with 26S protease inhibitor (Bortezomib or Bortz). Total proteins were extracted and separated on 6.5% SDS-PAGE and probed with anti-Myc and anti-RPT5 antibodies. RPT5 was used as loading control. (B) Bar graphs showing the quantitative PIF protein levels in those backgrounds from three biological replicates (n= 3). In each graph, PIF protein level in Col-0 background without Bortezomib treatment was set as 1. Error bars indicate mean ± SD. * indicates P < 0.05.

### Overexpression *PIFs* partially suppresses the cop phenotypes of *cop1-4* and *spaQ*

If the reduced level of PIFs in the *cop1-4* and *spaQ* backgrounds contributes to the cop phenotypes of these mutants, we hypothesized that an overexpression of these PIFs in the *cop1* and *spaQ* mutants is expected to suppress the cop phenotypes. To test this hypothesis, we overexpressed four PIFs (TAP-PIF1, PIF3-Myc, PIF4-Myc and PIF5-Myc) in the *cop1-4* and two PIFs (TAP-PIF1 and PIF5-Myc) in the *spaQ* backgrounds and examined their phenotypes in the dark. Results show that while the hypocotyl lengths of TAP-PIF1/cop1-4 and PIF3-Myc/cop1-4 are comparable to those of *cop1-4,* the hypocotyl lengths of PIF4-Myc/*cop1-4* and PIF5-Myc/*cop1-4* are significantly longer compared with *cop1-4* (Figure 3A-D). Moreover, all four PIF overexpression lines in the *cop1-4* mutant displayed significantly smaller cotyledon angle compared to that of *cop1-4,* suggesting that PIFs suppress the cop phenotypes of *cop1-4.* These data also suggest that the cop phenotype of *cop1-4* and *spaQ* are partially due to a reduced level of PIFs.

**Figure 3:**
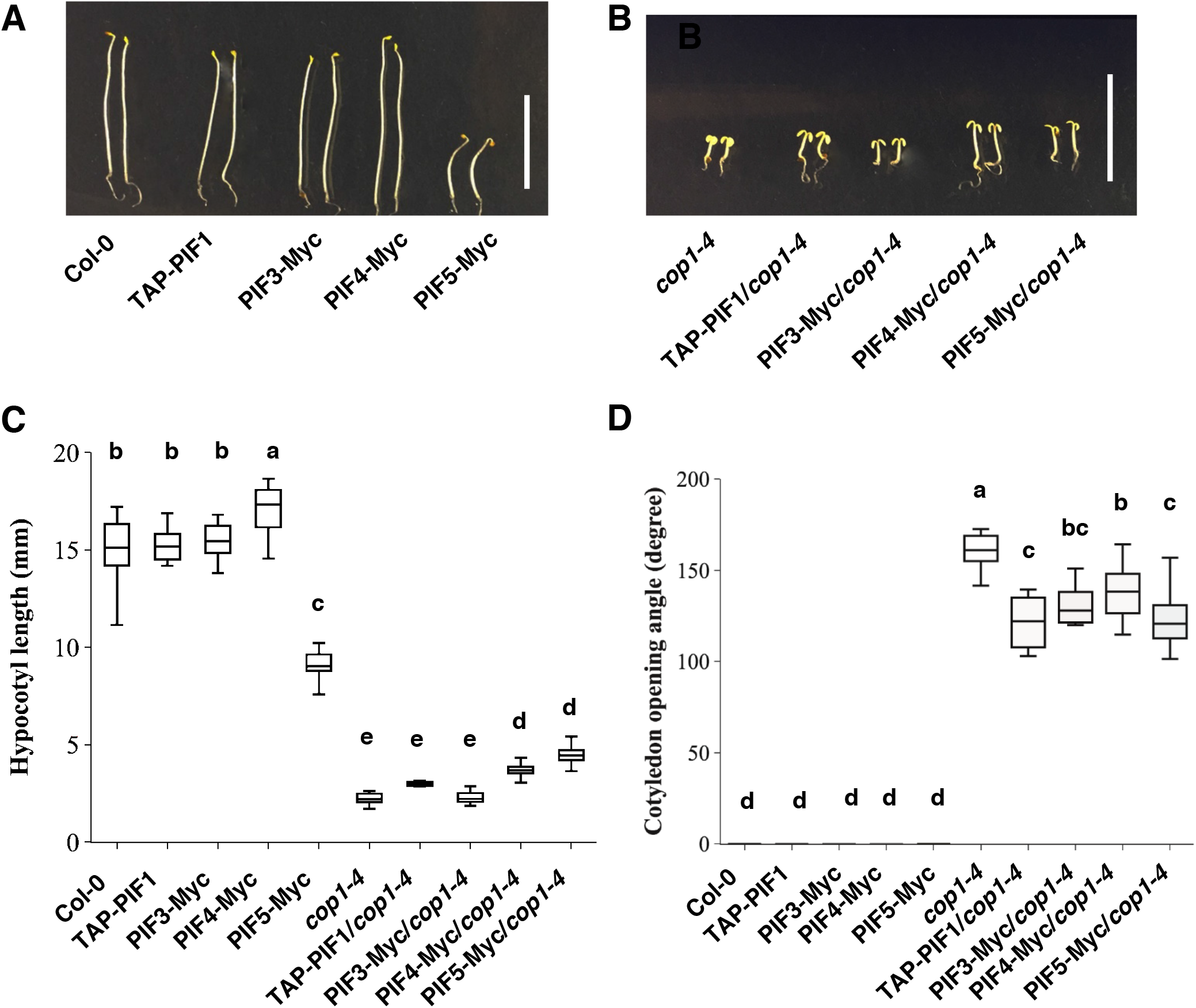
Overexpression PIFs partially suppresses the constitutive photomorphogenetic phenotypes of *cop1-4.* (A-B) Visible phenotypes of the 4-day-old dark-grown PIF overexpression in Col-0 and cop1-4 backgrounds. White bar = 10mm. (C) Box plots represent the hypocotyl lengths and cotyledon opening angle measurements. Three independent biological replicates were performed with an average of 30 PIF overexpression in Col-0 and cop1-4 seedlings grown under the same conditions as described above. Significant difference between different genotypes was determined using one-way ANOVA and Tukey’s HSD tests, indicated by different letters.

### *cop1-4, spaQ* and *pifQ* display a large overlapping set of co-regulated genes

Previously, the cop phenotypes of the *cop1* mutant was demonstrated to be mainly due to the high abundance of the positive regulators (e.g., HY5/HFR1/LAF1 and others) in the dark (Hoecker, 2017). Because PIFs are unstable in the *cop1* and *spaQ* mutants in the dark (Figure 1) (Bauer, Viczian, Kircher, *et al,* 2004; Leivar, Monte, Oka, *et al,* 2008; Zhu, Bu, Xu, *et al,* 2015), we hypothesized that COP1-, SPA- and PIF-regulated genes might overlap in the genome wide expression analyses. To test this hypothesis, we analyzed the results from a recent RNA-Seq experiment using the *cop1-4, spaQ* and *pifQ* mutant seedlings grown in darkness (Pham, *et al,* 2018a; Zhang, *et al,* 2013). Although these experiments were performed in two different laboratories and growth conditions were slightly different, the results show that a large proportion of the differentially expressed genes (1120) overlap among *cop1-4, spaQ* and *pifQ* (Figure 4A, B; Dataset S1). Approximately, 39% percent of the PIF-regulated genes display overlapping expression patterns with the COP1- and SPA-regulated genes. Among these 1120 genes, 483 genes are up-regulated and 431 genes are down-regulated in all three backgrounds compared to wild type (Figure 5A, B). Interestingly, 206 of the PIF-regulated genes display opposite regulation from the COP1- and SPA-regulated genes (Figure 4A, B; Dataset S3).

**Figure 4:**
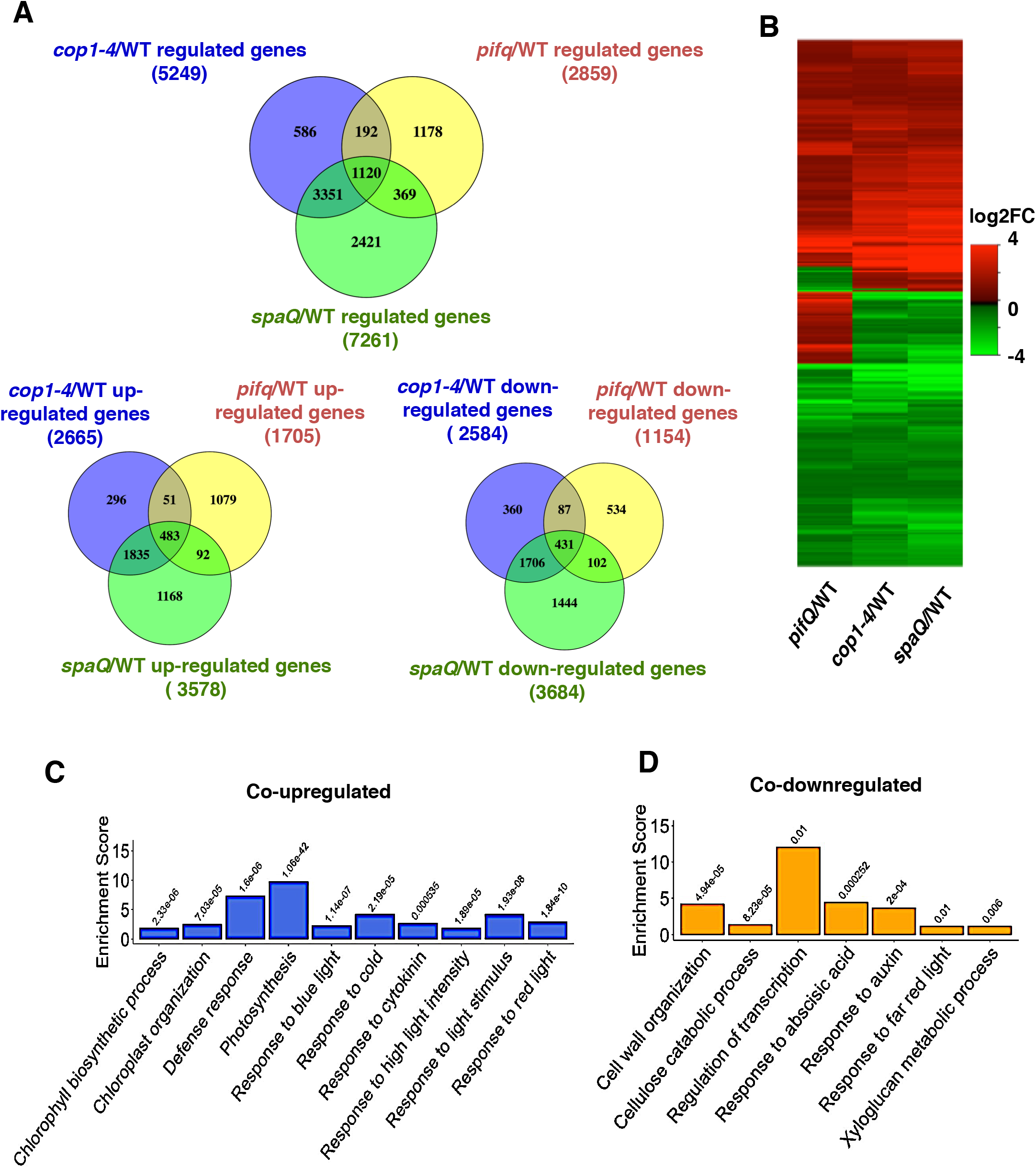
COP1 and SPA mediate light-regulated transcriptomic changes partly through PIFs. (A) Venn diagram showing 1120 co-regulated, 483 co-upregulated and 431 co-downregulated differential expressed genes in 3 different pairwise comparisons indicated *(cop1-4/WT, spaQ/WT* and *pifQ/WT).* (B)Hierarchical clustering displays 1120 DEGs in comparisons indicated. Data showing genes co-regulated identified as at least 2-fold difference in gene expression (FDR < 0.05) The color represents the log2 of fold change. *cop1-4/WT* and *spaQ/WT:* comparison of the expression profiles of dark-grown *cop1-4* and *spaQ* with Col-0, respectively. (C, D) Bar graphs showing Enrichment analysis of Gene Ontology Biological Processes significantly co-upregulated (C) and co-downregulated (D) in *cop1-4/WT, spaQ/WT,* and *pifQ/WT.* Enrichment scores indicate percentages, involved genes/total genes. Fisher Exact P-Values were presented on the top of each bar.

**Figure 5:**
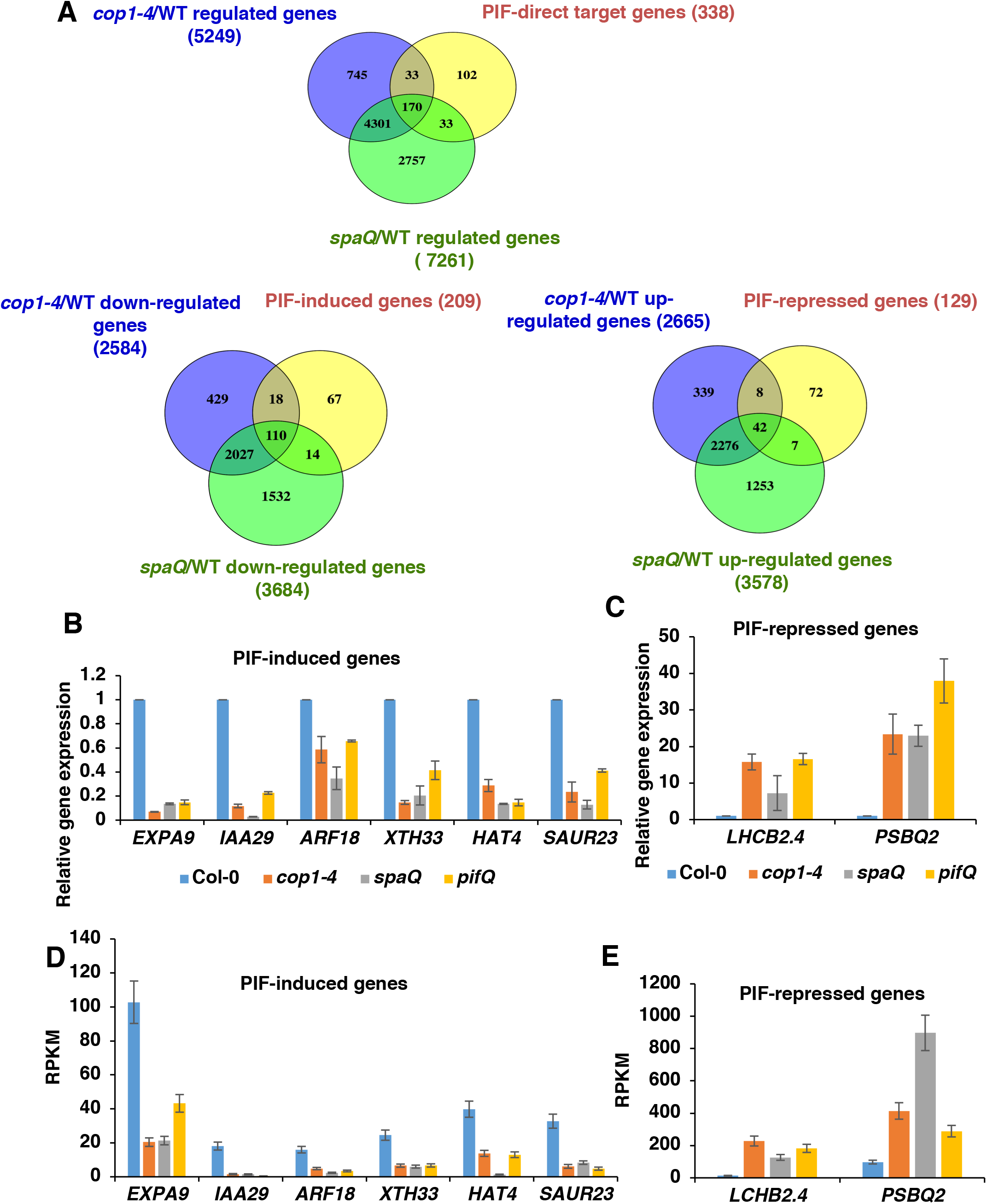
PIF-direct target genes are co-regulated in *cop1-4* and *spaQ* mutants. (A) Venn diagram showing among 338 PIF-direct target genes, 170 genes are coregulated by COP1 and SPA, 110 genes are down-regulated and 42 genes are up-regulated in *cop1-4* and *spaQ* mutants. (B) to (C) qRT-PCR shows the similar expression patterns of various PIF-induced genes (B) and PIF-repressed genes (C) in *cop1-4, spaQ* and *pifQ* in the dark. Gene expression level in mutants were normalized to *PP2A* and expression level in Col-0 was set as 1. (D) To (E) Expression patterns of various PIF-induced genes (D) and PIF-repressed genes (E) in *cop1-4, spaQ* and *pifQ* in the dark from RNA-seq data.

To identify the biological processes controlled by these co-regulated genes, we performed Gene Ontology (GO) analyses of the co-regulated genes and divided into two classes: up-regulated and down-regulated genes (Figure 4C, D; Dataset S2). A total of 94 enriched GO terms have been identified for these co-regulated genes. The co-upregulated genes are enriched in chlorophyll biosynthetic process, defense responses, photosynthesis, response to light stimulus (including red light and blue light), response to cold and cytokinin. The co-downregulated genes are involved in regulation of transcription, cell wall organization, response to hormones (abscisic acid and auxin), response to red light and also metabolic processes. These results are consistent with the cop phenotypes of these mutants.

Interestingly, 206 of the PIF-regulated genes display opposite regulation from the COP1-and SPA-regulated genes (Figure 4A, B; Dataset S3). GO analyses of the oppositely regulated genes (206 genes) between *pifQ* and *cop1-4/spaQ* using **D**atabase for **A**nnotation, **V**isualization and **I**ntegrated Discovery (**DAVID**) (Dataset S3) and GO Analysis Toolkit and Database for Agricultural Community (AgriGo) (Figure S4 and S5) show that PIFs function oppositely compared to COP1 and SPA proteins in a few biological processes. For example, many genes involved in responses to UV-B and flavonoid biosynthesis are down-regulated in *pifQ,* while they are up-regulated in *cop1-4* and *spaQ* (Figure S4; Dataset S3). Similarly, many other genes involved in defense responses, salicyclic acid (SA) metabolism and signaling are up-regulated in *pifQ,* while they are down-regulated in *cop1-4* and *spaQ* (Figure S5; Dataset S3. Thus, while PIFs and COP/SPA complex repress photomorphogenesis coordinately, they also function antagonistically in a few biological processes.

### Direct target genes of PIFs are co-regulated in *cop1-4* and *spaQ*

If COP1 and SPA repress photomorphogenesis in the dark in part by stabilizing PIFs, then the PIF direct target gene expression is expected to be affected in *cop1-4* and *spaQ.* Interestingly, among 338 PIF direct target genes (Pfeiffer, Shi, Tepperman, *et al,* 2014), 170 (>50%) genes are co-regulated by *cop1-4* and *spaQ* (Figure 5A; Dataset S4). Furthermore, among the 209 PIF-induced genes, 110 genes are down-regulated in *cop1-4* and *spaQ.* In addition, among 129 PIF-repressed genes, 42 genes are up-regulated in *cop1-4* and *spaQ* backgrounds (Figure 5A; Dataset S4). GO analyses revealed that the majority of these genes function in responses to red, far-red light signaling, auxin responses and regulation of transcription. Strikingly, the degree and direction of expression of these PIF direct genes are very similar among all three mutant groups.

To verify the RNAseq data by an independent method, we selected a subset of PIF direct target genes involved in auxin responses, cell wall organization and photosynthesis, and performed RT-qPCR analyses to determine the relative expression patterns in the *cop1-4, spaQ* and *pifQ* in comparison to wild-type. Results show a strikingly similar pattern among *cop1-4, spaQ* and *pifQ* for both PIF-induced and – repressed genes (Figure 5C, D), consistent with the RNAseq data (Figure 5D, E). These data suggest that the *cop* phenotype of the *cop1* and *spaQ* mutants might be partly due to the reduced level of PIFs and their target gene expression.

### HFR1 represses the transcriptional activity of PIF1

Previously, it was shown that HFR1, a HLH transcription factor is more abundant in the *cop1-4* and *spaQ* backgrounds compared to wild type (Hoecker, 2017). Because HFR1 inhibits the DNA binding activity of PIFs (Shi, *et al,* 2013), and the PIF levels are reduced in the *cop1-4* and *spaQ* (Figure 1 B-F; Figure S1A), we hypothesized that the high abundance of HFR1 in the *cop1-4* and *spaQ* backgrounds might contribute to the cop phenotypes of these mutants. To test this hypothesis, we selected a subset of the PIF1 direct target genes that are also known to be regulated by HFR1 and performed the RT-qPCR analyses from dark-grown seedlings of wild type Col-0, *pifQ, cop1-4* and *cop1-4 hfr1.* Results show that the selected genes are expressed at a reduced level in both *pifQ* and *cop1-4* mutant backgrounds similar to the RNAseq data (Figure 6A). Strikingly, the expression level of these genes is much higher in the *cop1-4 hfr1* double mutant background compared to *cop1-4* (Figure 6A). However, the increased expression of the PIF1 target genes in the *cop1-4 hfr1* might be either due to the high PIF1 protein level, and/or due to the loss of HFR1’s suppression of PIF1’s DNA binding activity. To distinguish between these possibilities, we performed the chromatin immunoprecipitation followed by quantitative PCR (ChIP-qPCR) for dark-grown seedlings expressing the TAP-PIF1 fusion protein in the *cop1-4* and *cop1-4 hfr1* backgrounds. We also examined the immunoprecipitated TAP-PIF1 protein level in all these backgrounds during the ChIP assay (Figure 6B, inset), and divided the promoter enrichments by the protein levels for each genotype to calculate the relative promoter occupancy of PIF1 independent of PIF1 protein level. Results show that the immunoprecipitated TAP-PIF1 protein level is lower in the *cop1-4* background, but higher in the *cop1-4 hfr1* double mutant background compared to the TAP-PIF1 only as previously reported (Figure 6B, inset) (Zhu, Bu, Xu, *et al,* 2015). The relative promoter occupancy of PIF1 show that the DNA binding activity of TAP-PIF1 was also reduced in the *cop1-4* background and increased in the *cop1-4 hfr1* background (Figure 6B). These data further extend the recent report that HFR1 suppresses PIF1’s DNA binding activity not only in imbibed seeds (Shi, Zhong, Mo, *et al,* 2013), but also in seedlings. Thus, HFR1 not only regulates the protein abundance but also the DNA binding activity of PIF1 in etiolated seedlings.

**Figure 6:**
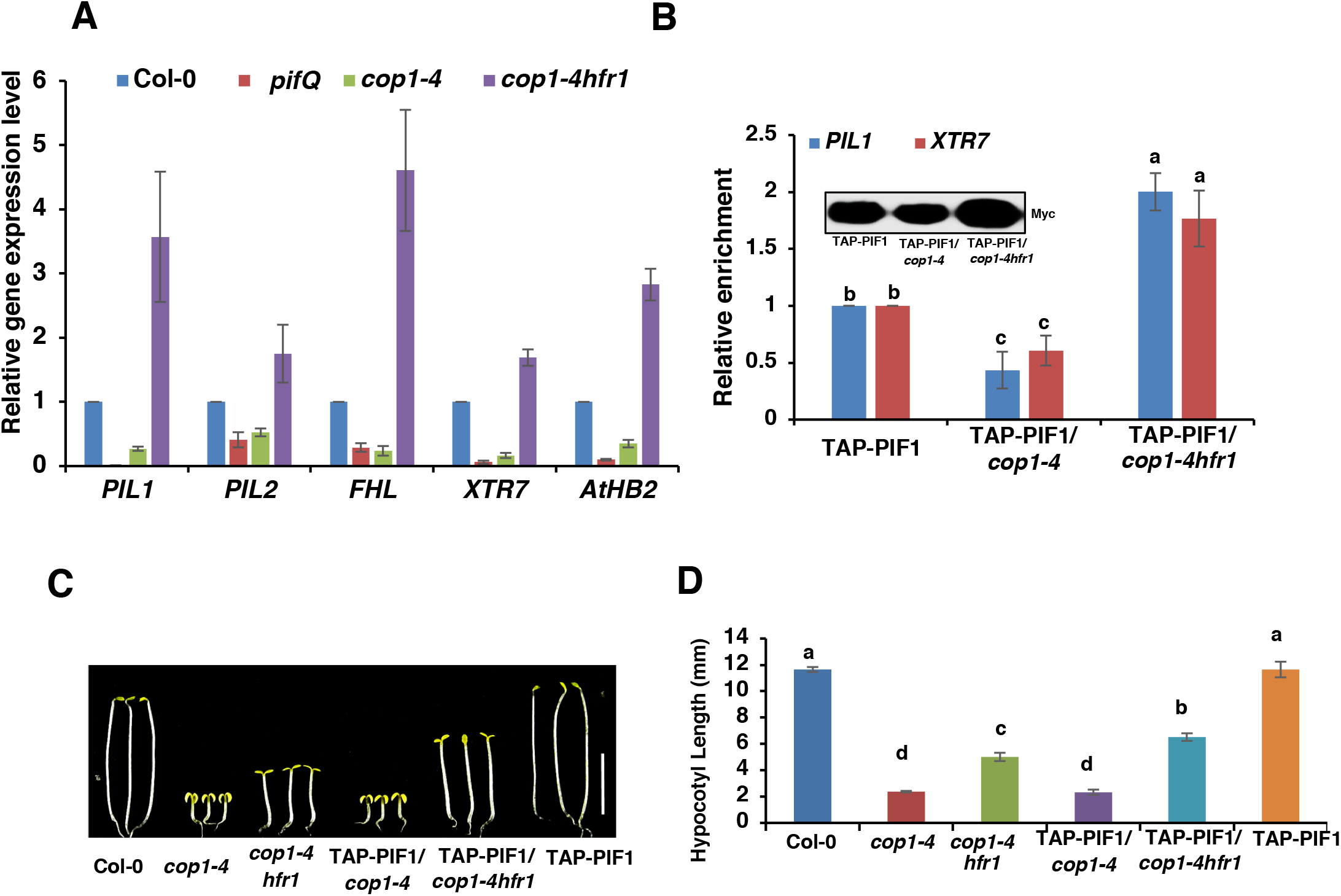
The transcriptional activation activity of PIFs is higher in *cop1-4hfr1* compared to *cop1-4* background. (A) The expression levels of PIF target genes are lower in the *pifQ* and *cop1-4* background, but are higher in *cop1-4 hfr1* background. Total seedling RNA was extracted from 3-day-old dark grown seedlings of wild type Col-0, *pifQ, cop1-4* and *cop1-4 hfr1. PP2A* was used as an internal control. Wild type Col-0 was set as 1. Error bars indicate standard deviation (n= 3 independent biological repeats). (B) The *PIL1* and *XTR7* promoter occupancies of TAP-PIF1 were reduced in *cop1-4* background but increased in *cop1-4hfr1* background. The ChIP-qPCR assays were performed on 3-day-old dark grown seeding expressing TAP-PIF1 fusion protein in *cop1-4* and *cop1-4hfr1* backgrounds. (C)Photographs of seedlings of wild type, *cop1-4, cop1-4hfr1, cop1-* 4/TAP-PIF1 and *cop1-4hfr1* /TAP-PIF1. Seedlings were grown in the dark for 5 days. White bar = 5 mm. (D) Bar graph showing hypocotyl lengths of various genotypes as described in (C). Error bars indicate standard deviation. Significant difference between different genotypes was determined using one-way ANOVA and Tukey’s HSD tests, indicated by different letters.

To examine the significance of regulation of PIF1 by HFR1, we measured the hypocotyl lengths for the dark-grown seedlings of wild-type, *cop1-4, cop1-4 hfr1, cop1-4/TAP-PIF1, cop1-4 hfr1/TAP-PIF1* and *TAP-PIF1.* Results show that the hypocotyl lengths of the *cop1-4 hfr1/TAP-PIF1* are much longer than that of the *cop1-4 hfr1* backgrounds (Figures 6C, D). The hypocotyl length of the *cop1-4* is similar to that of the *cop1-4/TAP-PIF1* possibly due to the reduced level of TAP-PIF1 protein level and/or increased sequestration of TAP-PIF1 by HFR1 in the *cop1-4* background. Thus, TAP-PIF1 has an increased function in regulating hypocotyl lengths in the *cop1-4 hfr1* background compared to only *cop1-4.*

## Discussion

Analyses of the cop mutants have played an important role in our understanding of the light signaling pathways in plants. The prevailing view of the molecular basis of the cop phenotype is that the increased abundance of the positively acting transcription factors (e.g., HY5/HFR1/LAF1 and others) in the *cop1-4* and *spaQ* mutants in the dark results in the cop phenotypes under darkness (Jang, Yang, Seo, *et al,* 2005; Osterlund, Hardtke, Wei, *et al,* 2000; Saijo, *et al,* 2003; Seo, Yang, Ishikawa, *et al,* 2003; Yang, *et al,* 2005a; Yang, Lin, Sullivan, *et al,* 2005b). Although several studies reported a reduced abundance of PIFs in various cop mutants compared to wild type (Bauer, Viczian, Kircher, *et al,* 2004; Leivar, Monte, Oka, *et al,* 2008; Ni, Xu, Tepperman, *et al,* 2014; Shen, Ling, Castillon, *et al,* 2008; Zhu, Bu, Xu, *et al,* 2015), the mechanism of this reduction and its contribution to the cop phenotype remain unknown. Here we show biochemical, molecular and genomic evidence supporting the hypothesis that the reduced PIF levels in the *cop1* and *spaQ* mutants contribute to their cop phenotypes. First, we show that PIFs are actively degraded in the dark in the *cop1-4* and *spaQ* backgrounds through the 26S proteasome pathway (Figures 1, S1). Second, the genome wide gene expression patterns largely overlap between COP1-, SPAs- and PIF-regulated genes with an altered expression of a set of PIF direct target genes (Figures 4, 5). Third, PIF1 is sequestered in the *cop1-4* background by increased abundance of HFR1 and possibly other HLH proteins, resulting in the reduced PIF activity in the *cop1-4* background (Figure 6). Fourth, an overexpression of PIF1 in the *cop1-4 hfr1* background promotes hypocotyl elongation in the dark. Fifth, an overexpression of four major *PIFs* in the *cop1-4* background suppresses the cop phenotypes of the *cop1-4* (Figure 3). Overall, these data strongly suggest that the reduction in the PIF levels and PIF activity in the *cop1-4* and *spaQ* backgrounds contribute to their cop phenotypes.

Despite similar morphological and molecular phenotypes among *cop1-4, spaQ* and *pifQ,* the GO analyses of the differentially expressed genes oppositely regulated between *pifQ* and *cop1-4/spaQ* reveals that these mutants also have distinct roles in plant signaling pathways. One of the striking differences is in the enrichment of the genes involved in the SA metabolism and signaling in *pifQ* compared to *cop1-4* and *spaQ,* suggesting that PIFs might suppress defense responses as previously discussed (Paik, *et al,* 2017). In fact, PIFs are known to promote growth possibly by suppressing the defense responses, as a trade-off between growth vs defense is a well-known phenomenon in plant growth and development (Paik, Kathare, Kim, *et al,* 2017). In contrast, the genes involved in the UV-B responses and flavonoid biosynthesis are down-regulated in *pifQ,* while they are upregulated in *cop1-4* and *spaQ.* Although, a role for PIFs in the UV-B signaling has not been examined in detail yet, a recent study implicated that PIFs might be involved in UV-B-induced leaf hyponasty (Fierro, *et al,* 2015). However, COP1 and SPA proteins function positively in the UV-B signaling pathways (Huang, *et al,* 2013; Tilbrook, *et al,* 2013). Overall, these analyses highlight both common and distinct functions of PIFs, COP1 and SPA proteins in regulating biological processes in plants.

One of the sources of the opposite functions between PIFs and COP1/SPA complex is the increased abundance of the positively acting transcription factors especially HFR1 in the *cop1-4* and *spaQ* backgrounds. HFR1 is an atypical bHLH protein that sequesters PIF activity as well as reduces the PIF abundance (Figure 6) (Hornitschek, *et al,* 2009; Shi, Zhong, Mo, *et al,* 2013; Xu, Kathare, Pham, *et al,* 2017). Similar to HFR1, HECATE family of bHLH proteins also inhibits PIF activity, and is degraded in the dark possibly by COP1/SPA complex (Zhu, *et al,* 2016). Thus, COP1/SPA complex might negatively regulate the abundance of factors that function antagonistically to PIFs.

In summary, we propose a revise model for the molecular bases of the cop phenotypes in plants (Figure 7). First, as previously hypothesized, an increased abundance of the positively acting transcription factors (e.g., HY5, LAF1, HFR1 and others) in the *cop1* and *spaQ* mutants promotes photomorphogenesis in the dark. Second, a reduced level of PIFs in the *cop1* and *spaQ* mutants contributes to the cop phenotype in the dark. Finally, a reduction in PIF activity due to the increased abundance of the atypical bHLH proteins (e.g., HFR1, HECATE and possibly others) in *cop1* and *spaQ* mutants additively promotes the cop phenotypes. It is notable that all three activities are tightly linked to each other contributing in concert to the skotomorphogenic and photomorphogenic developments in plants.

**Figure 7:**
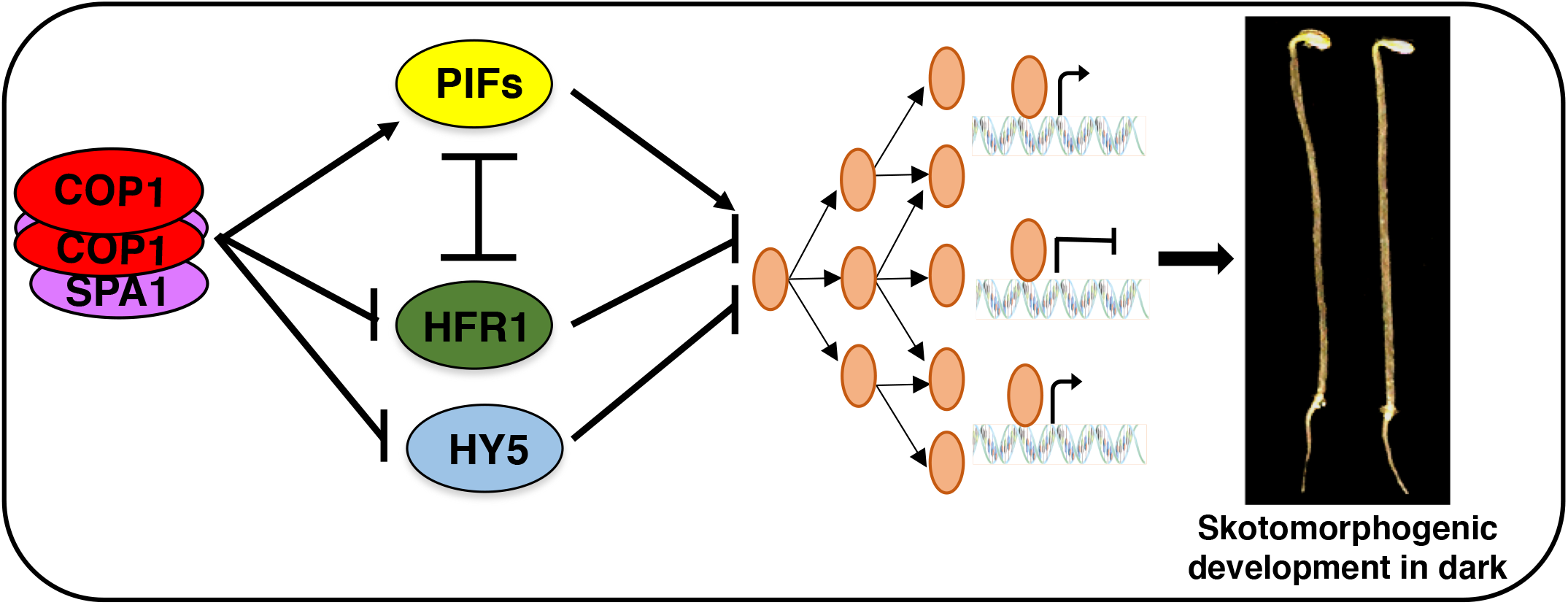
Model showing how COP1 and SPA proteins regulate various transcription factors to promote skotomorphogenesis in the dark. Mutation in COP1, SPA and PIFs results in constitutive photomophogenic (cop) phenotypes in the dark. PIFs and HFR1 reciprocally regulate their abundance, while HFR1 inhibits the PIF activity by sequestration. Regulation of these transcription factor abundance and activity by the COP1-SPA complex promotes skotomorphogenic development.

## Materials and Methods

### Plant materials, growth conditions and measurements

Columbia-0 (Col-0) ecotype of *Arabidopsis thaliana* seeds were used for all experiments. Seeds were surface sterilized and then plated on the Murashige and Skoog (MS) medium without sucrose. After stratified at 4°C for 3 days, seeds were exposed to white light for 3 hours at room temperature to trigger the seed germination before placing them back in the dark for an additional 4 days. These 4-day-old seedlings were used for protein extraction for Western blots, measurements of hypocotyl lengths and cotyledon opening angle phenotypes. A total of 90 seedlings from three biological replicates were measured for hypocotyl lengths and cotyledon opening angles using ImageJ software (http://rsb.info.nih.gov/ij/). Significant difference between different genotypes was determined using one-way ANOVA and Tukey’s HSD tests, indicated by different letters in the figures.

### Generation of transgenic lines

The *pif1, pifQ, cop1-4, cop1-4pif1,* cop1-4/TAP-PIF1 (Castillon, *et al,* 2009; Xu, *et al*, 2014; Zhu, Bu, Xu, *et al*, 2015), *TAP-PIF1* (Bu, Zhu, Yu, *et al*, 2011), *PIF3-Myc* (Park, *et al,* 2004), *PIF4-Myc* (Shor, *et al,* 2017) and *PIF5-Myc* (Sakuraba, *et al,* 2014) plants were previously published. *PIF* overexpression lines were crossed with *cop1-4* and *spaQ* mutants. The crossed homozygous lines were selected from F3 population using antibiotic selection. The *cop1-4* mutant was selected by sequencing. The *spaQ* homozygous lines were selected by genotyping *spa* mutants. Primers used for sequencing and genotyping are listed in Table S1. For generation of *cop1-4 hfr1*/TAP-PIF1, *cop-4 hfr1* was crossed into TAP-PIF1 to obtain F1 generation. Through genotyping and antibiotic selection (Gentamycin) of the F2 and F3 generation, we obtained *cop1-4 hfr1* /TAP-PIF1. Primers used for sequencing and genotyping are listed in Table S1.

### Transcriptomic analyses

To evaluate and compare the differential gene expression in *cop1-4* and *spaQ,* the raw read counts were obtained from our previous RNA-seq study (Pham, Hoecker and Huq, 2018a). Raw data and processed data for the total read counts of sequencing reads in Col-0, *cop1-4* and *spaQ* can be accessed from Gene Expression Omnibus database under accession number GSE112662.

RNA-seq was performed using 3-day-old dark-grown seedlings. Seeds were kept in the dark for 3 days at 4°C and exposed to 3hrs of while light. After 21h in the dark, plates were then treated with 2000 μmolm^-2^ far-red light for the true-dark condition as previously described (Leivar, Monte, Oka, *et al,* 2008). Total RNA was extracted after 2 days in the dark.

For the RNA-seq analysis, raw read quality was accessed using FastQC (http://www.bioinformatics.babraham.ac.uk/projects/fastqc/). Raw reads were then aligned to the *Arabidopsis* genome using Bowtie2 (Langmead and Salzberg, 2012) and TopHat (Trapnell, *et al,* 2012). The annotation of the *Arabidopsis* genome was obtained from TAIR10 (https://www.arabidopsis.org/). Read count data were performed by HTseq (Anders, *et al,* 2015) (http://htseq.readthedocs.io/en/master/index.html).

Differentially expressed genes in *cop1-4/WT* and *spaQ/WT* were identified using DESeq2 package (Love, *et al*, 2014). The differential gene expression was defined as those differ by > 2-fold with adjusted *P* value (FDR) < 0.05.

Differentially expressed genes in *pifQ* and PIF differential direct-target genes list were obtained from RNA-seq and ChIP-Seq data, respectively, under the accession number GSE43286 (Pfeiffer, Shi, Tepperman, *et al,* 2014). Venn diagrams were generated using Venny 2.1.0 (http://bioinfogp.cnb.csic.es/tools/venny/). Heatmaps were generated using DESeq2 and ComplexHeatmap package (Gu, *et al,* 2016) in the R statistical program. Gene Ontology (GO) enrichment analyses were performed using **D**atabase for **A**nnotation, **V**isualization and **I**ntegrated **D**iscovery (**DAVID**) v6.8 https://david.ncifcrf.gov/. GO bar graphs were generated based on the significant enriched terms with the lowest p value and FDR (< 0.05) for GO terms. Hieratical graph results for the GO term analysis of *cop1-4, spaQ* and *pifQ* regulated genes were also performed by GO Analysis Toolkit and Database for Agricultural Community (AgriGo http://bioinfo.cau.edu.cn/agriGO/index.php).

### Quantitative RT-PCR assay

For determining the transcript levels of *PIFs* and PIF-direct target genes by RT-qPCR assays, total RNA was extracted from the seedlings grown under the same conditions used for the RNA-seq experiment. One μg of total RNA treated with on-column DNase I (Sigma Aldrich, St. Louis, MO) was reversed transcribed using M-MLV reverse transcriptase (Thermo Fisher Scientific, Bartlesville, OK). RT-qPCR assay was performed using Power SYBR^®^ green (Applied Biosystems, Foster City, CA). Gene specific primers are listed in Table S1. *PP2A* (At1g13320) was used as the internal control to normalize the expression of different genes. The calculation of the levels of expression of different gene relative to *PP2A* is as follows: 2^ΔCt^, where ΔCt = Ct *(PP2A)*-Ct (specific gene) and Ct indicates the cycle threshold values. Relative expression was quantified from three biological replicates. Error bars indicate mean ± SD. Student’s t-test assuming unequal variances was performed, *P* values are indicated in each figure.

### Protein extraction and immunoblot analyses

For examination of COP1 protein level, 0.2g tissue from 4-day-old dark-grown seedlings was extracted in extraction buffer as described (Zhu, Bu, Xu, *et al,* 2015). Total protein was separated on 8% SDS-polyacrylamide gel electrophoresis (PAGE). Proteins were transferred to PVDF membrane and Western blots were detected with anti-COP1 or anti-RPT5 (Enzo Life Sciences, Farmingdale, NY) antibodies for endogenous COP1 and RPT5, respectively.

For PIF protein levels in Col-0 and mutant backgrounds, total protein from 50 seedlings was extracted using 50μl urea extraction buffer (48% urea (w/v), 0.1M phosphate buffer pH 6.8, 10mM Tris-Cl pH 6.8, 1 mM phenylmethylsulfonyl fluoride (PMSF) and 1× protease inhibitor cocktail). Samples were centrifuged at 16,000g for 10 min and heated 65°C for 10 min. Supernatants were analyzed on 6.5% SDS-PAGE gels and detected using anti-Myc (dilution 1/1000, Catalog number OP10-200UG, EMD Millipore, MA) and anti-RPT5 antibodies (dilution 1/3000, Catalog number: BML-fPW8245-0100, Enzo Life Sciences, Farmingdale, NY).

For treatment of 26S proteasome inhibitor (Bortezomib, catalog number B-1408, LC Laboratories), 4-day-old dark-grown seedlings were transferred to 5 mL liquid MS media containing 40 μM Bortezomib and incubated in the dark for 4 hrs. Total protein was then extracted using the urea extraction buffer as described above. For quantitation of protein levels, ImageJ software was used to measure the band intensities from three independent biological replicates, and normalized to RPT5 protein levels. Error bars indicate mean ± SD. Student’s t-test assuming unequal variances was performed, *P* values are indicated in each figure.

### Chromatin immunoprecipitation (ChIP) assay

The ChIP-qPCR assays were performed on 3-day-old dark-grown seedlings expressing TAP-PIF1 fusion protein in *cop1-4* and *cop1-4hfr1* backgrounds. Anti-Myc (rabbit) antibody was used to immunoprecipitate TAP-PIF1 and associated DNA. DNA was amplified using primers specific to the G-box fragment or control regions. Anti-Myc (mouse) antibody was used to determine the immunoprecipitated TAP-PIF1 protein level in each background. Both the TAP-PIF1 promoter enrichment from the ChIP-qPCR and TAP-PIF1 protein level quantified by ImageJ were set as 1. The relative enrichment of the fold-change of the *cop1-4*/TAP-PIF1 and *cop1-4hfr1*/TAP-PIF1 were first normalized compared with the TAP-PIF1 only for their promoter enrichment and protein levels, respectively, and then divided the promoter enrichments by the protein levels for each repeat. Final averages of three independent biological repeats for each genotype were calculated and shown as the bar graph. One biological repeat of the TAP-PIF1 protein level was shown as an example in inset. Error bars indicate standard deviation (n= 3 independent biological repeats).

## Acknowledgments

We thank Drs. Xing Wang Deng for sharing *cop1* mutant, Ute Hoecker for sharing *spaQ* mutant, Giltsu Choi for sharing *PIF3-Myc* and *PIF5-Myc* seeds and Woe Yeon Kim for COP1 antibody. We thank the Huq lab members for the technical support and critical reading of the manuscript.

## Author contributions

V.N.P., X.X. and E.H. designed experiments. V.N.P., and X.X. carried out experiments. V.N.P., X.X. and E.H. analyzed data and interpreted the results. V.N.P. wrote the article. V.N.P., X.X. and E.H. edited the manuscript.

## Supplementary material

Supplementary material is available online.

